# Species abundance plays a dominant role in local and regional extinctions in freshwater communities and allows the identification of selective extinctions

**DOI:** 10.1101/219469

**Authors:** Ryan J. Almeida, Elizabeth G. Biro, Lauren M. Woods, Kevin G. Smith

## Abstract

Recent declines in global biodiversity emphasize that understanding the factors that determine extinction risk should be a priority for ecologists and conservation biologists. A key question is whether extinctions are nonrandom and selective, in which case knowledge of selectivity may help predict and prevent future extinction. We suggest, however, that a premature focus on the identification of selective, trait-based determinants of extinctions risk is problematic if the potential importance of stochastic extinction processes are not first considered. Within this context we aimed to determine the roles that stochastic extinction and species abundance play in extinction risk by applying a rarefaction-based null model approach to analyzing biodiversity declines and extinctions in an experimental system. We focused on aquatic macroinvertebrate declines and extinction caused by predation by fish (*Lepomis cyanellus*) in semi-natural freshwater mesocosms. We found that null-predicted local extirpations based on the random loss of individuals were a significant predictor of observed local extirpations, and that the majority of observed extinctions were consistent with stochastic mechanisms of extinction, as predicted by a rarefaction model. We were able to identify a number of selective extinctions that were not predicted by the rarefaction model, and while these were relatively rare, they contributed to greater-than-expected loss of diversity at both local (mesocosm) and regional (whole experiment) spatial scales. Our results confirm that species abundance and occupancy are among the most important factors in identifying extinction risk in response to a disturbance. Moreover, owing to our use of a stochastic null model, we also conclude that measures of abundance are important indicators of extinction probability because they are operated on by the random loss of individuals, suggesting that stochastic extinction is an important process in this system and in biodiversity loss in general.

## Introduction

A focal point of modern ecology and conservation is the unsustainably high rate of species loss that has occurred in recent history as human population growth, development, and anthropogenic global climate change have contributed to local, regional, and global declines in global biodiversity (Dirzo & Raven 2003; Barnosky *et al*. 2011; Steffen *et al*. 2011). As an ecological process, extinction is a fundamental determinant of global patterns of biodiversity, both throughout geological history and in modern times and understanding extinction is critical to understanding biodiversity. Furthermore, as a focus of conservation biology, understanding extinction bias, i.e., why some species are more prone to extinction than others, is essential to the effective conservation and management of biodiversity. This focus on extinction selectivity and bias is conceptually related to the role played by extinction in macroevolution (Raup 1994), in which extinction is considered part of the selective process leading to the “survival of the fittest”. Indeed, research often emphasizes extinction as a nonrandom process and extinction selectivity and bias are appropriate frequent topics of study in ecology and conservation (McKinney 1995; McKinney & Lockwood 1999; Peters 2008; Jonathan L. Payne *et al*. 2016). Yet when comparing populations of the same species, it is not surprising that smaller populations are more likely to be lost in face of a disturbance than larger populations (Harrison 1991; Husband & Barrett 1998; Melbourne & Hastings 2008). This phenomenon suggests that while species-specific extinction selectivity plays a role in species loss, random processes based on population size may also contribute significantly to patterns of extinction. In general the balance between random processes and nonrandom processes in determining extinction risk is rarely studied and the relative importance of each process is often unclear. Ultimately, additional work focusing on whether extinction processes are primarily selective and based on species traits, or are primarily stochastic and therefore based strictly on species abundance is essential for understanding and successfully predicting extinctions.

Previous studies focusing on the mechanisms of species extinction rarely directly address the contribution of stochastic processes to extinction risk. Instead, the importance of species functional traits and deterministic filtering are highlighted (Russell *et al*. 1998; Purvis *et al*. 2000; Cardillo *et al*. 2005; Bielby *et al*. 2006). While functional or phylogenetic selectivity clearly contribute to the extinction process, it has long been recognized that stochastic extinction processes are also important, particularly in small, isolated, or declining populations (Shaffer 1981; Wilcox & Murphy 1985; Pounds *et al*. 1997). And yet the role played by random probability is rarely taken into account in the analysis of extinction patterns. For reasons of simple probability, we follow others in suggesting that stochasticity operates upon abundance and rarity to determine extinction probabilities (Shaffer 1981; Simberloff 1986; Raup 1992; Pounds *et al*. 1997) and that there is predictive merit to explicitly accounting for the stochastic effects of abundance on extinction probability when analyzing patterns of extinction (Wilcox & Murphy 1985; McKinney 1997; Smith *et al*. 2009).

Our suggestion that patterns of extinctions are reflective of underlying random processes follows from the potential for large differences in extinction rates among taxa, even when species are neutral, with the only differences among species being abundance or range size. As a simplified heuristic example, we generated a simulated community of 30 species drawn from a log series (Fisher 1934) with *x* = 0.90 to create a realistic abundance distribution. We then randomly removed 50% of the individuals from the population in a rarefaction procedure (Gotelli & Graves 1996), simulating a significant habitat loss event or other large loss of individuals from a single location. The results from this simulation (Fig. 1) are not unexpected, but emphasize the expectation of dramatic differences in extinction probability among taxa even when the loss of individuals is strictly random and species are neutral. While these results are expected based on probability, such stark differences in extinction rates among species could be inferred as resulting from differences in species fitness or specific traits. In such a case, if this role of abundance and random probably is not first taken into account, a random pattern of extinction could easily lead to potentially erroneous conclusions regarding extinction selectivity and risk among taxa.

**Figure 1.**
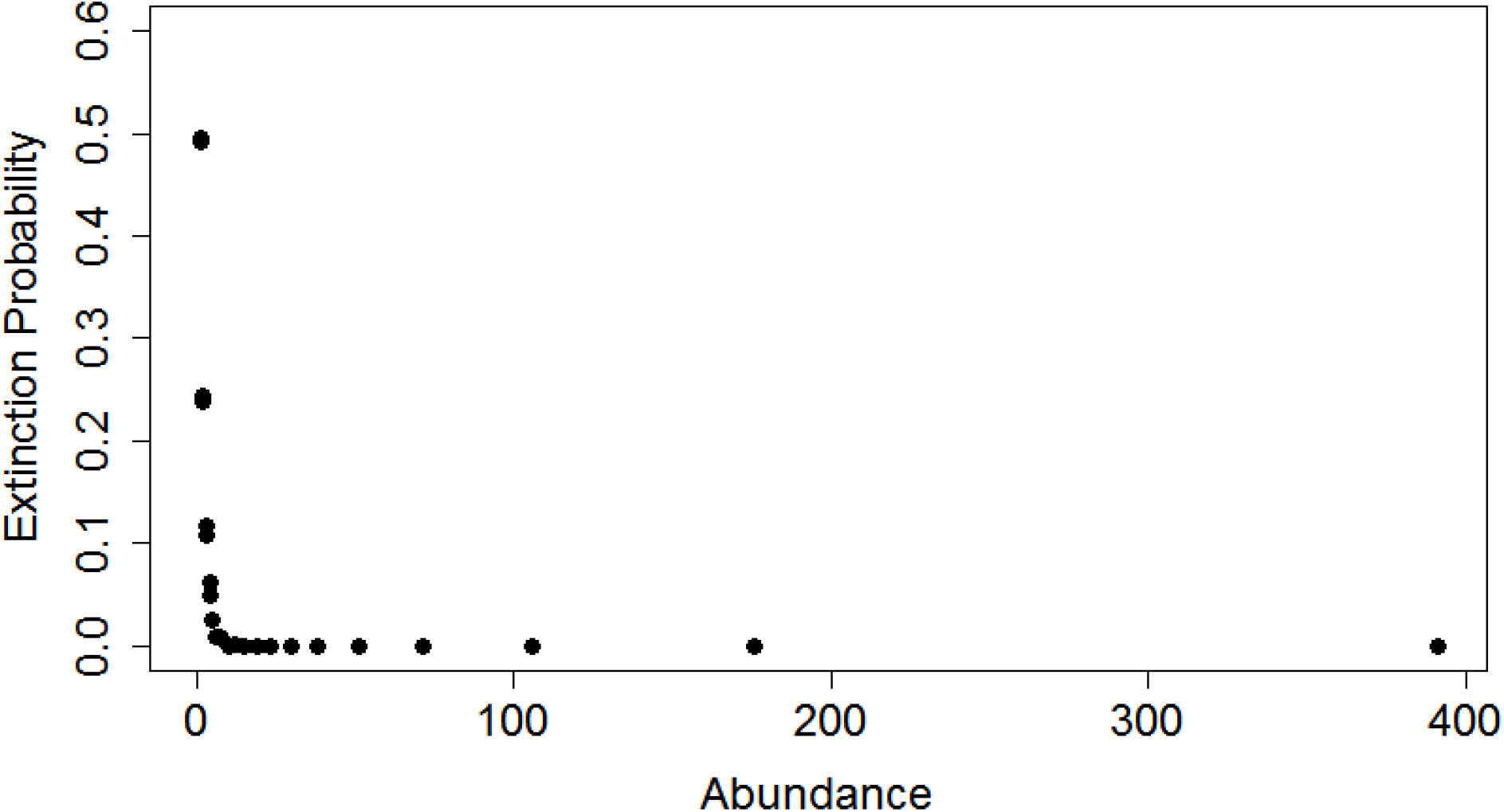
Relationship between species extinction probability and abundance, based on simulated data for a community of 30 species drawn from a log series. Each point represents the null-expected extinction probability for a species based on its abundance alone. Extinction probabilities were calculated via rarefaction, simulating the effects of a random loss of 50% of the individuals in the original community.

Within this context, we used a stochastically neutral (*sensu* Vellend *et al*. 2014) null model approach based on rarefaction (Gotelli & Graves 1996) to analyze local and regional extinctions with the goal of examining the role that abundance and the stochastic loss of individuals play in determining species extinction risk. We apply this approach to data from a mesocosm experiment in which predatory green sunfish (*Lepomis cyanellus*) were added as a disturbance, leading to local (within-mesocosm) and regional (among-mesocosm, within-treatment) extinctions of macroinvertebrates. Our study consisted of artificial ponds to which macroinvertebrates were added and allowed to colonize naturally prior to the introduction of sunfish. Our null model approach allows us to determine whether stochastic processes were a significant predictor of extinction risk by modeling expected species loss based on the random removal of individuals from the community. In contrast, if fish predation is dominated by selective processes based on species functional traits and irrespective of abundance, then we hypothesize that our null model will have limited predictive ability in determining which species go extinct post-disturbance. Our results suggest that even with putatively selective predators (Chase *et al*., 2009), stochastic processes act via the random loss of individuals form local communities and play an important and potentially predictive role in the local and regional extinction process. This strong signal of stochasticity adds nuance to the perspective that species extinctions are selective, deterministic events that may reliably be predicted by species traits.

## Methods

### Mesocosm establishment

This study took place at Washington University’s Tyson Research Center in Eureka, MO. In 2007 we established 32 1100-L stock tanks, to which we added ca. 5 cm of topsoil. We filled tanks with local well water, and inoculated them with aquatic submerged and emergent plants. We also added aliquots of concentrated zooplankton mixtures from a series of local ponds. All tanks were left open to the environment to allow for colonization by organisms from the local species pool. We added several gastropod species that would not be able to naturally colonize each tank. These mesocosms were used in other experiments in 2008 and 2009, but then were left fallow until this study commenced in 2012. In May and June of 2012 we surveyed all tanks for their existing aquatic biodiversity, which we then supplemented with macroinvertebrates collected from four local ponds and added to a random subset of half of the mesocosms (Table S3). After these initial surveys, we covered tanks with fiberglass mesh. In early June 2012 we introduced 3-5 green sunfish (*Lepomis cyanellus*) to 16 of 32 mesocosms ensuring at least one male and one female was introduced to each. We stratified the fish introductions across treatments from the past studies in 2008 and 2009 to ensure that legacies from prior treatments did not affect the present study. Control mesocosms (those to which fish were not added) were used to identify the local extinctions that were not associated with the addition of fish. These species were removed from analysis of species losses caused by fish (see *Data Preparation*, below).

### Biodiversity sampling

We sampled all aquatic macroinvertebrates in each mesocosm via area-constrained surveys in which two 25 cm-diameter cylinders were placed in each mesocosm and sampled exhaustively for living macroinvertebrates using aquarium nets (“chimney sampling”, as described in Chase *et al*., 2009; Woods *et al*., 2016). All animals were held temporarily to allow for live identification by an experienced researcher (author EGB) with reference to standard field guides and keys. We separately collected organisms living on the inside walls of each mesocosm using four sweeps of an aquarium net (8 x 10 cm) along each inside wall of a mesocosm. Finally, after the above biodiversity sampling we observed each tank individually (5 minutes per mesocosm and 10 sweeps of a D-net) for the presence of rare species that were not recorded using other methods. These rare species were recorded as having one individual. After identification, we returned all animals to their original mesocosms to ensure that sampling did not affect the abundance or extinction probability of taxa in our study.

We sampled each mesocosm using these methods prior to the introduction of fish, which occurred on 8 June 2012. We conducted two post-treatment biodiversity surveys, three and six weeks after the introduction of fish (2 July and 24 July, respectively).

### Null Model

We used a null model approach to compare observed declines in abundance and observed extinctions in our experiment to expected declines (Gotelli & Graves 1996). We constructed a null model in R (R Core Development Team, 2017) that applies a rarefaction approach to species loss (Fig. 2). Species richness is often a function of total individual abundance at a particular site (Srivastava & Lawton 1998) and rarefaction determines if observed differences in species richness are strictly the result of differences in abundance or sample size (Sanders 1968). As typically applied, rarefaction produces expected species richness values based on the repeated subsampling of individuals from a larger sample (Simberloff 1972; Gotelli & Graves 1996; Gotelli 2001; Gotelli & Colwell 2001) and our model operates under these same premises. Specifically, for a given decline to *x* individuals at a particular site (in our study, a mesocosm), the null model randomly samples *x* individuals from the pre-decline dataset in 1000 simulated declines such that the pre-decline communities are randomly rarefied down to the lower-abundance post-decline communities. Observed and expected (i.e., rarefied) species richnesses or diversities are then compared.

**Figure 2.**
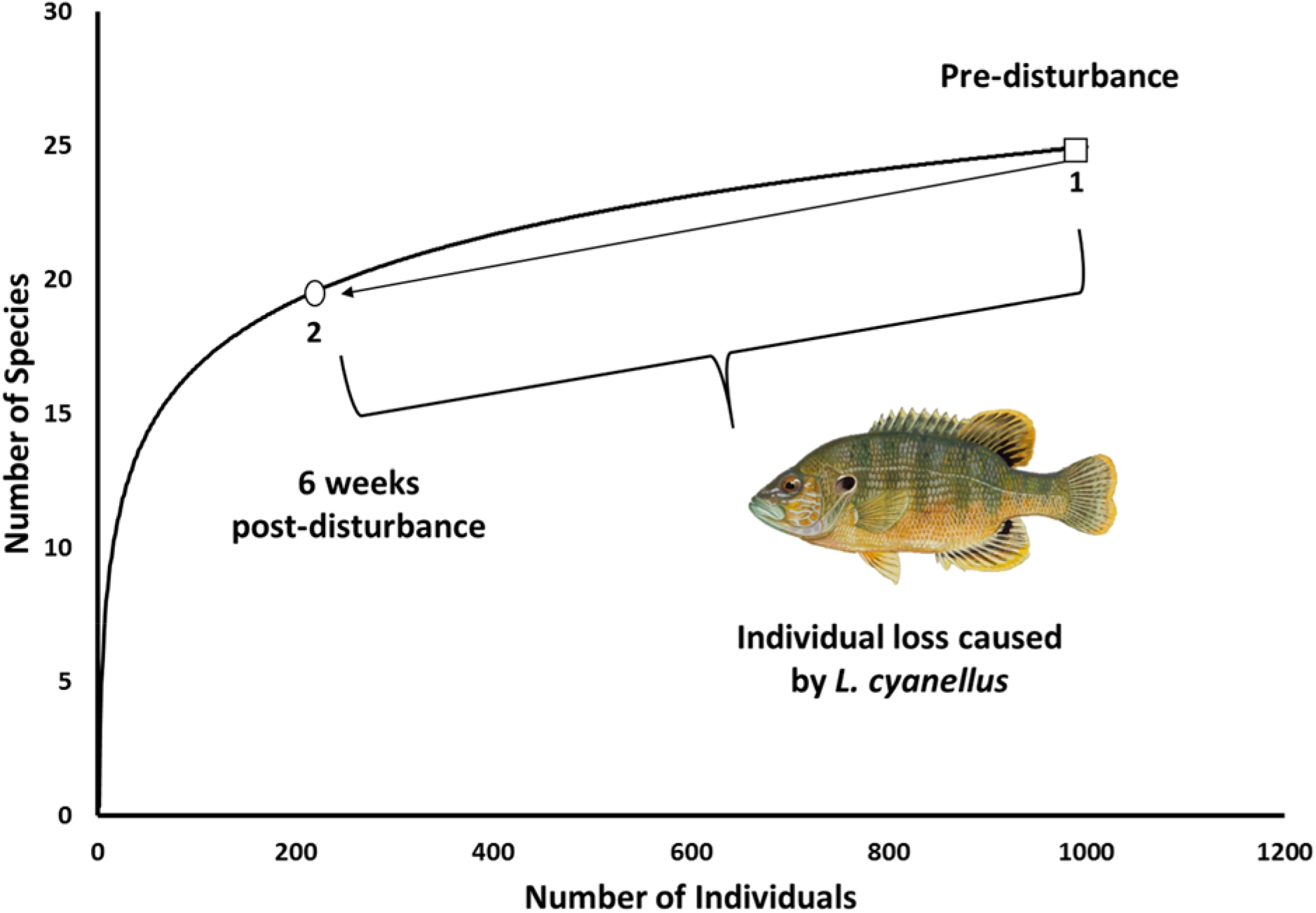
**Graphical representation of rarefaction as a null model for extinction**. Under the assumptions of the null model, species loss is dictated solely by the random removal of individuals. Box 1 shows the species accumulation of a pre-disturbance community, while Box 2 presents an expectation for species richness, and therefore species loss, assuming neutrally stochastic extinction processes, in this case as caused by fish predation. *L. cyanellus* drawing by Ruane Daver (public domain).

We expand on the typical output of rarefaction simulations (rarefied species richness) by further calculating expected abundances for individual species post-decline and expected random probabilities of extinction for each species post-decline. We define the random probability of local extinction, PeL(*i*) for each species *i* as the proportion of rarefaction runs that result in an abundance of zero (local extinction) for species *i*. These simulations were conducted simultaneously for each site within a treatment “region” of multiple mesocosms (originally 16, but see below), which allowed us to also calculate P_eR_(*i*), the random probability of regional extinction from all sites in the treatment “region”. We then used these P_e_ values to assess whether observed extinctions deviated from null expectations; the loss of a species with P_e_ ≤ 0.05 was considered an unexpected extinction and evidence of deterministic or selective extinction processes. Extinctions that could not be distinguished statistically from null expectations (P_e_ > 0.05) were considered consistent with extinctions predicted by neutrally stochastic processes and the purely random loss of individuals.

### Data Preparation

Data requirements of our rarefaction-based null model required the following modifications to our biodiversity data from each sampling period. First, we excluded from our analyses a single mesocosm in the fish treatment, owing to fish extinction in this mesocosm. We also removed from the analysis ten species that were not present prior to the introduction of fish but appeared in later surveys. Because each site was closed to immigration to prevent the introduction of new species throughout the sampling period, the appearance of a species after the introduction of fish was likely due to the transition from undetectable life stages (e.g., eggs or planktonic larvae). Second, some species may have declined or disappeared from mesocosms in our study owing to phenology (e.g., metamorphosis of aquatic larval odonates into aerial adults) rather than mortality from predation. Many aquatic macroinvertebrates undergo an ontogenetic habitat shift; once the juveniles have metamorphosed to the adult life stage, they leave the site to search for new breeding ground. As a result, the loss of individuals due to phenology could lead to an over-estimate of the effects of fish predation. To address this issue, we analyzed each species’ decline in both control (non-fish) and treatment (fish) sites. If a species experienced large declines in control as well as fish treatments, we considered those losses to have been caused by phenology and therefore removed those species from the analysis. We provide a complete list of removed species in Supplementary Table S1.

### Linear Regression

To assess the relative importance of stochastic processes during a local extinction event, we compared observed extinctions to the mean number of simulated extinctions at each individual site. We conducted linear regression analysis to determine if simulated (expected) extinctions were a significant predictor of observed extinctions. If extinctions were perfectly predicted by neutral stochastic processes, we would expect a 1:1 relationship to exist between observed extinctions and expected extinctions (i.e., observed extinctions could not be distinguished from null model predictions.) Alternatively, a lack of relationship between observed and expected extinctions would suggest the presence of strong selective processes that lead to local and regional extinctions occurring irrespective of species abundance.

## Results

### Loss of Biodiversity

After data preparation, 28 macroinvertebrate species occurred across all sites. As expected, the addition of green sunfish led to large reductions in macroinvertebrate diversity and abundance. On average, fish treatment sites experienced an 81.3% (± 10.1% SD) average reduction in individuals in each mesocosm, corresponding to an average of 5.8 (± 2.67 SD) local extinctions after six weeks of exposure to sunfish predation. Fish predation accounted for a larger loss of individuals than occurred naturally in the controls.

The rarefaction analysis provided predictions for both the number of expected extinctions, and probabilities of extinction for each individual species. Of the observed local extinctions in the post-fish survey, an average of 5.8 extinctions were expected based on random processes alone (Table 1). We observed similar patterns with regional extinctions. At the regional level, eight extinctions had occurred six weeks after fish were added, with four of these extinctions being consistent with stochastic extinction based on rarefaction.

**Table 1.** **Mean (± 95% CI) expected and unexpected local extinctions after introduction of sunfish**. Mean extinctions are averages of extinctions from all treatment sites (n= 15). Unexpected extinctions were defined as extinctions of species with a P_eL_ ≤ 0.05.

| | Mean Extinctions with $P_{eL} > 0.05$ | Mean Extinctions with $P_{eL} \leq 0.05$ | Mean local extinctions (observed) |
| --- | --- | --- | --- |
| Post-Fish (Six Weeks) | $5.8 \pm 1.50$ | $0.72 \pm 0.36$ | $5.8 \pm 1.35$ |

### Relationship between expected and observed extinctions

Of the 15 analyzed tanks treated with sunfish, 14 sites had a larger number of observed local extinctions than would be expected. Linear regression analysis demonstrated a strong significant relationship between expected extinctions and observed extinctions (Fig. 3, F_1,13_ = 32.62 p = 0.0001, R^2^ = 0.72).

**Figure 3.**
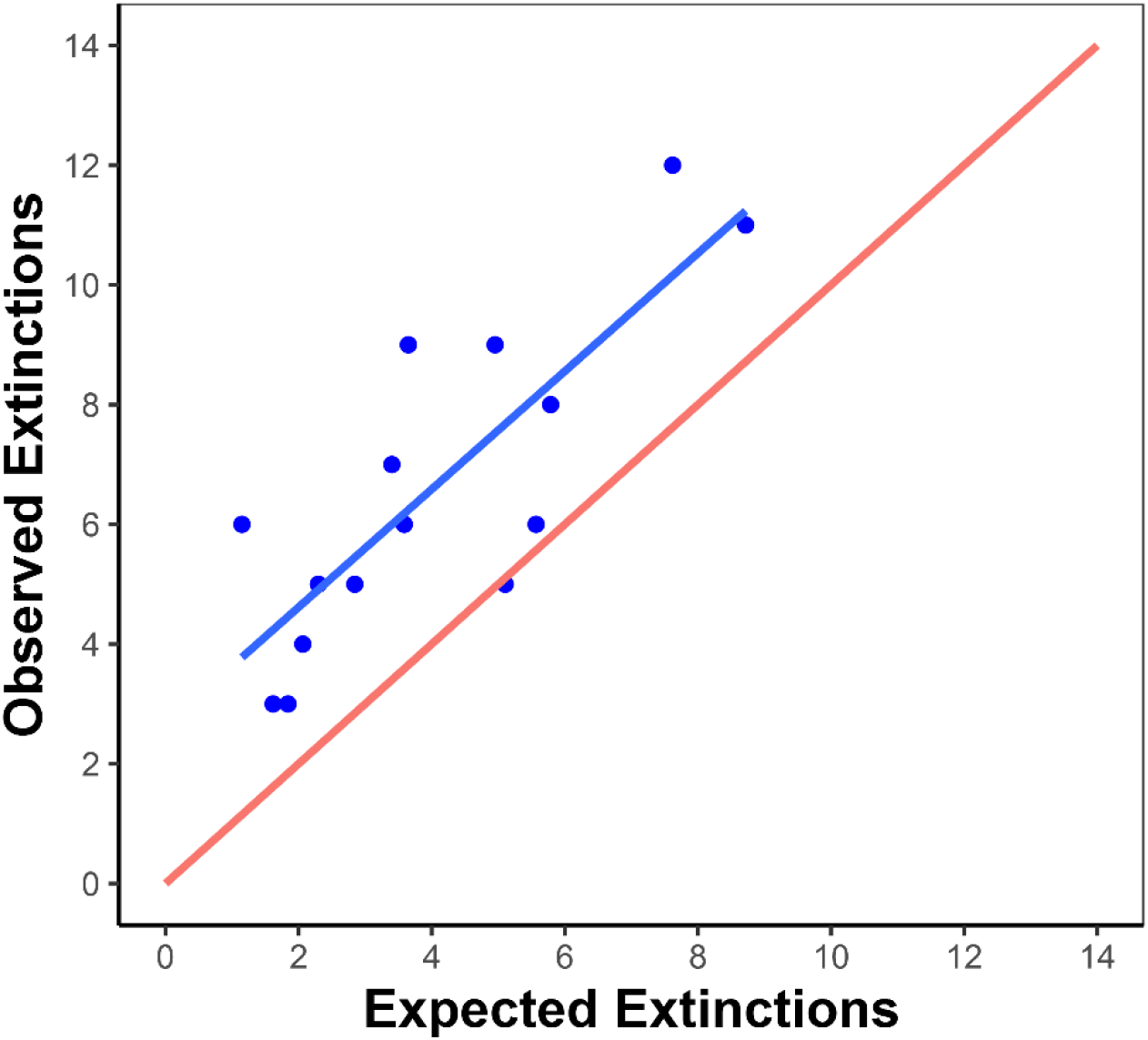
**Expected vs. observed local extinctions after second post-fish survey**. Each point represents a tank treated with sunfish (n = 15), with expected extinctions defined as the mean number of simulated extinctions at that site after three weeks of fish predation. The red line indicates expected relationship if expected extinctions were a perfect predictor of observed extinctions (slope = 1). Expected extinctions were a significant predictor of observed extinctions over this time scale (Linear Regression, F_1,13_ = 32.62, p = 0.0001, R^2^ = 0.72).

### Spatial Scaling

To determine if the dominance of stochastic processes was consistent across spatial scales, we compared expected and observed local extinctions to regional extinctions. In the post-fish survey, there were significantly more observed local extinctions than expected extinctions (Fig. 4; T-test; t = −2.55, df = 26.59, p = 0.017). This pattern persisted at the regional level, as there were significantly more observed regional extinctions than predicted by the null model (Fig. 4). The null model predicted 3.05 (95% confidence interval 1, 5) regional extinctions, while we recorded 9 observed regional extinctions.

**Figure 4.**
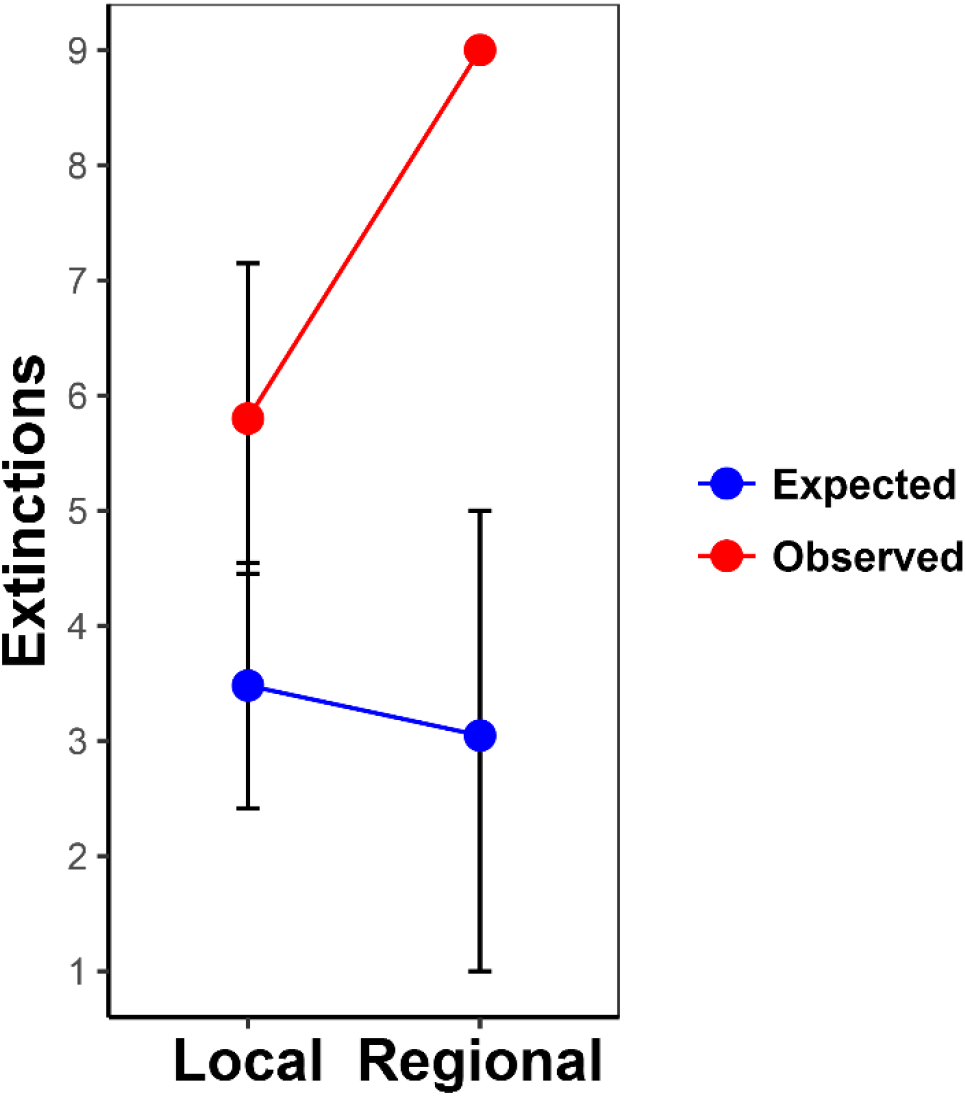
**Scaling of local and regional species richness in treatment sites after second post-fish survey**. Expected local extinctions were the mean number of extinctions simulated by the null model at each site, while expected regional extinctions were the mean number of times a species was simulated to go extinct at fifteen analyzed sites. 95% confidence intervals are shown when applicable. There were significantly more observed extinctions than expected extinctions at both the local and regional spatial scales.

## Discussion

The goal of our study was to clarify the role species abundance and stochastic extinction processes play in biodiversity loss in a model system of predatory fish in aquatic freshwater communities. We used a novel application of a standard ecological method, rarefaction, to create a null expectation of species richness and abundance based on the random removal of individuals. If disturbances result in the random loss of individuals, then species abundance will be the sole predictor of extirpation and extinction and rarefaction should reliably predict the number of extinctions and even the extinction of individual species. This approach allowed us to explore the importance of stochastic extinction by assigning each individual species an extinction risk (P_e_) based on abundance and random chance alone. Our analysis produces a number of insights into the relative importance of stochasticity in the extinction process. In particular, although the number of observed extinctions was usually greater than the number predicted based on rarefaction, we found that the number of randomly-expected extinctions was a significant and important predictor of the number of observed extinctions after predators were introduced. This provides evidence for the predictive importance of abundance to extinction risk and further emphasizes the dominance of stochastic processes in local and regional species extinctions. In sum, rarefied species richness explained 68% of the observed variation in number of local extinctions.

Despite this evidence for the importance of stochastic extinctions, we found that most mesocosms experienced more extinction than predicted based on random loss via individual rarefaction. These differences, while modest at the local scale, were enhanced at the regional scale (Fig. 4). Combined with the fact that expected stochastic extinctions proved to be a significant predictor of observed extinctions, we interpret this result as potentially indicative of an additive effect of selective processes on local species loss. At each site, while stochastic processes largely determine which species go extinct (Table 1), a smaller number of additional, unpredicted extinctions of abundant species occurred leading to almost uniformly greater loss of biodiversity than predicted by the null model (Fig. 3 & 4). These nonrandom extinctions may be attributed to a number of factors. For instance, fish may have been exhibiting partially selective predatory behavior by targeting and/or avoiding particular species, in addition to feeding opportunistically and randomly on individuals as they were encountered. The mechanisms for such selective predation are potentially diverse. For example, previous studies focusing on selective predation by fish have proposed that increased visibility and body size make prey more susceptible to predation (Zaret & Kerfoot 1975; Bakker *et al*. 1997; Green & Côté 2014) while other traits including physical and chemical defenses make prey less susceptible to fish predation (Kats & Dill 1998; Kolar & Wahl 1998; Chappell & Smith 2016), although several of these patterns may be more strongly correlated with prey abundance. In our study, the backswimmer *Buenoa* and the aquatic beetles of *Tropisternus* are two examples of species that were driven extinct regionally (from all mesocosms) despite having a PeR < 0.05, strongly suggesting selective extinction owing to fish predation. Backswimmers forage in the water column and frequently occur close to the surface, which puts them at heightened risk of fish predation (Cook & Streams 1984). Additionally, species of *Tropisternus*, which is a widespread and relatively abundant beetle genus, exhibit strong aversion to selecting natural habitats that are inhabited by predatory fish, suggesting that this taxon may frequently be targeted by fish (Resetarits 2001). Thus, the selective predation of preferred macroinvertebrate species that were locally abundant in our study could have led to the additional observed selective extirpations (see also Supplementary Materials, Table S2).

The magnitude of regional extinction in our experiment was consistently greater than predicted by the rarefaction-based null model. This result is consistent with the scale-dependence of patterns of extinction and biodiversity loss, where losses of low-occupancy species accumulate among sites such that greater loss of diversity is seen at the regional level than would be predicted locally (Chase *et al*. 2009; Smith *et al*. 2009). For each species in our experiment, a regional population ranged from anywhere between one and 16 local populations, with different abundances within each local population. Because abundance could vary greatly across local populations, the null model highlights how selectivity manifests itself most predominantly at the regional level. Locally, it may be difficult to distinguish whether an extinction is the product of demographic stochasticity or trait-based filtering, especially for species of low abundances, for which stochastic extinction probabilities (P_eL_) are high. At the regional level, however, selective extinctions are easier to identify given the low probability (P_eR_) that even the least abundant species will be randomly lost from every site in which it occurs. Thus, patterns of selective extinction will be more easily identified at the regional level (Smith *et al*. 2009). In fact, 67% of unexpected regional extinctions (2 out of 3: *Laccophilus* and *Tropisternus*, Table S2) in the fish treatment were of species expected to be extirpated at the local level based on low species abundances. This highlights the outcome that while locally rare species may be buffered by widespread populations, widespread disturbances may still result in selective extinctions that would not be expected randomly (Smith *et al*., 2009). Given that there are more rare species in populations than there are common species (Magurran 2004), being able to differentiate the relative magnitudes of stochastic and deterministic processes and understand how they are affected by spatial scale is critical to understanding the factors behind extinction risk. This scale-dependency of biodiversity is a documented phenomenon (Chase & Leibold 2002; Hartley & Kunin 2003; Smith *et al*. 2009) that has important implications for conservation, as understanding regional extinction risk is analogous to understanding the factors driving the loss of global biodiversity.

The above selective extinctions could only be documented in our study because our null model is abundance-based. Although they come from an experimental system, these findings have important implications to criticisms of current conservation policy, which have called for more preventive measures to “keep common species common”, rather than solely focusing on rare and threatened taxa (Grenyer *et al*. 2006; Gaston 2010; Cardillo & Meijaard 2012). Specifically, these criticisms often warn against “conservation complacency” (Lindenmayer *et al*. 2011), where the susceptibility of common species to large declines goes unnoticed due to a narrow conservation focus on the protection of rare species. These declines in common species are occurring and have been observed over the past 20 years (León-Cortés *et al*. 1999; Gaston & Fuller 2007; Inger *et al*. 2015), yet such discoveries are infrequently reflected in conservation policy (Gaston & Fuller 2008; Lindenmayer *et al*. 2011). We suggest that our approach, which allows researchers to identify unexpected declines and losses of widespread and abundant species, can make a significant contribution to the formulation of more robust and inclusive conservation efforts.

Our study presents a different approach to the analysis of local and regional extinction processes at a time when the global extinction crisis is a focus of ecological and policy discourse. When participating in this discourse, it is imperative that conservation biologists have a complete understanding of extinction mechanisms to formulate accurate and effective predictions and policy recommendations. Past research has addressed the interplay of stochastic and selective processes in patterns of species diversity (Chase 2007; Chase *et al*. 2009; Farnon Ellwood *et al*. 2009; Smith *et al*. 2009; Swenson *et al*. 2011; Segre *et al*. 2014) through focusing on changes in β diversity, often based on species incidence alone. However these approaches are limited by ignoring the potentially important role abundance plays in determining community structure (Tucker *et al*. 2016) and local/regional extinction probability (this study). Our results add empirical evidence, from a controlled experiment, that species abundance, and by extension stochastic processes, play an important role in determining extinction risk at local scales. The fact that our results come from a manipulative experiment is important, as a critical limitation in extinction biology is that analyses of past *in situ* extinctions are necessarily correlative and cannot be replicated. In contrast, an experimental approach allows for the replication of the extinction process, thereby allowing for a more rigorous examination of extinction not only in model species, but in relatively high diversity model systems.

Finally, while our approach is trait-agnostic and therefore deliberately ignores differences among species, we do not dispute the importance of trait-by-environment interactions in the local extinction process. Species abundances, occupancies, and range sizes emerge in part from interactions between species traits and the environment (Rabinowitz 1981) with extinctions resulting from severe trait-environment mismatches. However, if a detailed knowledge of species functional traits is required to explain past extinctions and predict future extinctions, then most species in the tree of life cannot possibly benefit from species-specific conservation action. In contrast to this requirement for detailed knowledge of species traits, our results suggest that measures of species abundance alone may provide a valuable starting point for the anticipation of biodiversity losses. In summary, our findings suggest that knowledge of species diversity, abundance, and the potential magnitude of disturbance may allow for surprisingly informative minimum predictions of how many species—and which species—may be at risk of local and regional extinction.

## Acknowledgements

Muxi Yang and Sarah Jacobs assisted with field work for this project, including setting up the experiment and collecting data. Tyson Research Center (Washington University in St. Louis) provided space and logistical support for this project. Financial support for this project was provided by Washington University in St. Louis, Davidson College, and the National Science Foundation through grants to KGS (DEB 0816113 and DEB 1650554).

